# Regulation of the trade-off between cold stress and growth by *BcGSTF10*

**DOI:** 10.1101/2023.08.03.551777

**Authors:** Yunlou Shen, Guangpeng Wang, Jiajun Ran, Yiran Li, Huiyu Wang, Qiang Ding, Ying Li, Xilin Hou

**Affiliations:** National Key Laboratory of Crop Genetics & Germplasm Innovation and Utilization, Key Laboratory of Biology and Genetic Improvement of Horticultural Crops (East China), Ministry of Agriculture and Rural Affairs of China, Engineering Research Center of Germplasm Enhancement and Utilization of Horticultural Crops, Ministry of Education of China, Nanjing Agricultural University, Nanjing 210095, China; Nanjing Suman Plasma Engineering Research Institute Co.,Ltd. Nanjing 211162, China

**Keywords:** Cold stress, plant growth, BcGSTF10, BcICE1, BcCBF2, NHCC.

## Abstract

Cold stress represents a significant threat to global agricultural productivity and food security. Typically, plant resistance to cold stress is accompanied by a growth deficit and a reduction in yield. In this study, we have discovered a novel gene, *BcGSTF10*, which has not been previously reported to be involved in low-temperature stress. On the one hand, biochemical and genetic analyses have demonstrated that BcGSTF10 interacts with BcICE1 to promote the expression of *CBF* genes, thereby enhancing freezing stress tolerance in non-heading Chinese cabbage [NHCC, *Brassica campestris (syn. Brassica rapa)* ssp. *chinensis*] and *Arabidopsis*. On the other hand, the expression of the *BcGSTF10* gene is directly repressed by BcCBF2, while *BcGSTF10* exhibits a positive role in growth in both NHCC and *Arabidopsis*. This dual function of *BcGSTF10* indicates its pivotal role in balancing low-temperature stress and growth, which will inform the development of strategies to breed climate-resilient and high-yield crops, thereby contributing to sustainable agriculture.

## Introduction

Approximately 64% of the terrestrial land regions on Earth experience an average minimum temperature below 0℃ (Rihan et al., 2017). As sessile organisms, plants are more susceptible to cold stress than free-roaming animals (Zhu JK.,2016). Low-temperature stress can exert negative impacts on multiple physiological traits of plants such as growth, development, geographical distribution, and overall crop yield at a global level (Pearce., 2001, Ding et al., 2019). Therefore, throughout the process of long-term evolution, plants have developed complex regulatory mechanisms to balance the relationship between low temperatures and growth. Plants accumulate proline, sugars, and antifreeze proteins at the physiological level to withstand cold stress. These substances enhance the stability of the cell membrane, reduce membrane damage, improve cytoplasmic osmotic pressure, minimize the formation of intracellular ice crystals, and remove reactive oxygen species (Kidokoro et al., 2022, Dong et al., 2009). At the molecular level, an increasing number of regulatory pathways have been discovered for understanding the response mechanism of cold stress, including signaling molecules, protein kinases, and transcription factors. The most widely recognized regulatory pathway is the ICE1-CBFs-COR pathway (Stockinger et al., 1997; Liu et al.,1998; Shi, Y.,2015; Ding et al.,2018). The expression of COR (cold-regulated) genes has been proven to be essential for cold stress in plants (Gilmour et al.,1998). DREB1s/CBFs respond to low-temperature stress by binding directly to the promoter of COR genes. In this pathway, CBF gene induction occurs earlier than that of COR genes, while the cold-induced ICE acts upstream of CBFs (Chinnusamy et al.,2003). ICE1, as an important positive regulator of the CBF gene, has been extensively studied for its post-translational modification and regulation. OST1 (OPEN STOMATA1) is a protein kinase, which phosphorylates ICE1 under low-temperature signals, inhibits its interaction with E3 ubiquitin ligase HOS1, enhances the transcriptional activity of ICE1, and thereby improves the cold resistance (Ding et al., 2015). Phosphorylation of MPK3/MPK6 destabilizes ICE1 and negatively regulates the expression of CBFs, thereby affecting the cold tolerance of plants (Li et al., 2017a). BIN2 interacts with and phosphorylates ICE1, which leads to the facilitation of the interaction between ICE1 and HOS1, and consequently accelerates the HOS1-mediated degradation of ICE1, ultimately resulting in the negative regulation of cold tolerance in plants (Ye et al., 2019). These evidences demonstrate the central role of ICE1 in the low-temperature pathway. Interestingly, the ICE1 gene, as a constitutively expressed gene, is not induced by low temperature at the transcriptional level. Therefore, investigating how the low-temperature signal is transmitted to ICE1 and further affects plant cold tolerance is of great significance.

Plant glutathione transferases (GSTs) are a significant group of multifunctional enzymes that are divided into four classes (theta, zeta, tau, and phi) (Lan et al., 2009; Liu et al., 2015; Marrs et al.,1997). They play extensive roles in several aspects such as plant development, endogenous metabolism, stress tolerance, and xenobiotic detoxification (Nianiou-Obeidat et al., 2017). Overexpression of glutathione-s-transferase in tobacco plants results in changes in GPOX activity, oxidation-reduction status, and the induction of anti-oxidant mechanisms, ultimately leading to increased tolerance of tobacco seedlings to salt and low-temperature stresses (Roxas et al., 2000). The natural variation in the *OsGSTZ2* allele is responsible for the cold sensitivity in rice (Kim et al., 2011). Under various abiotic stresses, the expression of *OsGSTL2* is induced. Overexpression of the *OsGSTL2* gene in rice enhances its tolerance to drought, salinity, and cold stress (Kumar et al., 2013a; Kumar et al., 2013b). Although there have been many reports showing that GSTs are involved in both biotic and abiotic stress, little is still known about the endogenous substrates of plant GSTs and the mechanisms of how their activities are regulated.

Despite significant progress in elucidating the molecular mechanisms underlying low-temperature stress regulation over the past 20 years, the balance between stress tolerance and crop productivity remains a significant challenge in future crop engineering and agricultural applications (Feng et al.,2020). Early research indicated that low temperatures could affect the distribution of auxin, leading to stunted plant growth (Morris.,1979; Wyatt et al., 2002; Nadella et al.,2006; Shibasaki et al. 2009). Previous research has demonstrated that *Arabidopsis* can protect its stem cell niche from chilling stress by utilizing a selective cell death mechanism that is regulated by both auxin and DNA damage responses (Hong et al., 2017). Overexpression of *CBFs* resulted in abnormal plant growth and significantly delayed flowering time compared to the wild type (Jaglo-Ottosen et al., 1998; Gilmour et al., 2000; Gilmour et al., 2004; Park et al., 2015). Under low-temperature conditions, *cbf1/cbf2/cbf3* triple gene mutants were significantly larger than those of the wild type (Jia et al., 2016). The overexpression of *CBF1* in Arabidopsis thaliana caused a significant upregulation of the GA-inactivating GA2-oxidase gene, thereby resulting in a notable reduction in levels of bioactive gibberellin (GA). Consistently, the stunted growth and delayed flowering characteristics observed in plants overexpressing *CBF1* were rescued by application of exogenous gibberellin (GA). These results suggest that the *CBF1* gene may participate in suppressing the endogenous GA content of plants as part of balance plant growth and chilling stress (Achard et al., 2008).

Non-heading Chinese cabbage [NHCC, *Brassica campestris* (syn. *Brassica rapa*) ssp. *chinensis*] is favored by consumers for its high yield and nutritional value (Li et al.,2022). Nevertheless, the adverse effects of cold stress extend to both yield and quality. Although research on cold tolerance in agricultural plants is increasing recent years, study of non-heading Chinese cabbage is relatively scarce. In this study, through Y2H yeast screening, we have identified *BcGSTF10*, which encodes a glutathione-s-transferase. In previous studies, this gene was considered incapable of responding to low temperature in *Arabidopsis* (Ryu et al.2009). Nevertheless, we found that this gene plays an important role in balancing cold stress and yield in non-heading Chinese cabbage. *BcGSTF10* overexpression enhanced cold tolerance and promoted crops yield. Further research found that low-temperature signals are transmitted to *BcGSTF10*, thereby activating the transcriptional activity of BcICE1. However, prolonged low temperature inhibits the expression of *BcGSTF10*, thereby hindering crop growth. The findings have unveiled the means through which crops regulate their growth and manage cold stress, hence presenting novel strategies to enhance their productivity under low-temperature conditions.

## Materials and methods

### Plant materials and growth conditions

*Arabidopsis thaliana*, *Nicotiana benthamiana* and NHCC plants were grown at a constant temperature of 21℃±2°C, under a long-day photoperiod consisting of 16 hours of light and 8 hours of darkness, with a light intensity of 150 µmol m^-2^ s^-1^.

### Plasmid construction and genetic transformation

The coding sequences of *BcGSTF10* and *CBF2* were amplified from the cDNA of NHCC. To generate overexpression vectors, full-length *BcGSTF10*, *BcCBF2* were cloned into the pRI101 and pCAMBIA1300-GFP plasmids, respectively. The NHCC transformation and generation of transgenic lines were carried out using our previously established procedures (Guo et al., 2021) and were also subjected to identification (Supplementary Fig. 2A, B). Transient overexpression assay in NHCC leaves were determined as previous described (Wu et al., 2022). The BcGSTF10-OE, BcCBF2-OE lines were obtained by transforming 35S:BcGSTF10 and 35S:BcCBF2 construct into wild type Arabidopsis thaliana and identifying (Clough and Bent., 1998) (Supplementary Fig. 2C, D). The BcGSTF10 overexpression line was crossed with the BcCBF2 overexpression line, resulting in the generation of BcGSTF10-OE BcCBF2-OE lines, which were subsequently identified (Supplementary Fig. 2E). The specific primers used are listed in Supplemental Table 1.

### Cold tolerance assays

For *Arabidopsis thaliana*, the experiment on freezing tolerance reported earlier has undergone minor adjustments (Ye et al., 2019). *Arabidopsis* seedlings were grown under normal conditions for 12-14 days and subjected to a freezing assay in a freezing chamber. The freezing treatment starts at 1°C and decreases by 1°C per hour until it reaches -6°C. After recovery, seedlings that could still grow new leaves were scored as surviving seedlings. Four-leaf stage NHCC were used for cold stress, the seedlings were placed in a freezing incubator and incubated at -1°C for 24 hours, then transferred to a normal growth incubator for a 12-hour recovery period. Samples were taken and photographed for subsequent experiments.

### Ion Leakage Assays

Electrolyte leakage assays were performed as described by Li et al. (2017b) as follows, collect the frozen leaves and place them in tubes containing 8 mL of deionized water. Shake them for 20 minutes at 22℃. Then, their electrical conductivities (EC) were measured, giving S1. Next, the samples underwent a 30min boiling process, followed by 30min of vigorous shaking at a temperature of 22℃. The EC value of the samples labeled as S2. The EC of deionized water was set as S0. Ion leakage was calculated as: (S1 −S0)/(S2 – S0).

### DAB and NBT staining

Rosette leaves from *Arabidopsis* and NHCC were collected, both before and after undergoing low-temperature treatment, to be used for DAB and NBT staining. The leaves were immersed in DAB buffer (1 mg/mL) or NBT buffer (0.5 mg/mL) and shaken at 100 rpm in a dark environment. After 8 hours, the DAB or NBT buffer were replaced with bleaching solution (consisting of ethanol, acetic acid, and glycerol in a ratio of 3:1:1), and boiled at 95°C for 30min to remove the pigments. Finally, color changes in the leaves were observed and documented with images.

### Y2H assays and screening assays

Y2H assay was performed according to protocol (Clontech, cat. No. 630489). Since BcICE1 self-activates, we generated pGBD-BcICE1-F2 by inserting the truncated BcICE1 fragment (254-446 aa) into pGBD. We employed pGBD-BcICE1-F2 as a bait to screen for interacting proteins in the NHCC cDNA library. The ORFs of BcGSTF10 and truncated BcICE1 fragments were inserted into pGBD and pGAD vectors, respectively. The recombinant plasmids were co-transformed into yeast strain Y2H Gold and plated on medium lacking Trp and Leu (SD/-Trp-Leu) at 28°C. The colonies were then shifted to SD/-Trp-Leu-His-Ade medium for interaction screening. The primer pairs used for gene cloning are listed in Supplementary Table 1.

### BiFC assays

The full length CDS sequence of *BcICE1* and *BcGSTF10* were cloned into the 35S-pSPYNE-YFP^N^ and 35S-pSPYCE-YFP^C^ vectors, respectively. *Nicotiana benthamiana* leaves were co-infected with the transformed *Agrobacterium* mixture. YFP fluorescence was observed with a confocal laser-scanning microscope (Zeiss LSM 510 Meta, Jena, Germany). The primers used in this experiment are shown in Supplementary Table 1.

### Pull-down assays

Full-length BcGSTF10 and BcICE1 were recombined into pCold-TF and pMAl-c2X vector, respectively. The His, His-BcICE1 and MBP-BcGSTF10 proteins were expressed from *E. coli Rosetta* (Tolo Biotech, Shanghai, China) with isopropyl-β-d-thiogalactoside (IPTG) (Beyotime, Shanghai, China). Pull-down assays were carried out utilizing the HIS-tagged Protein purification Kit (Beyotime, Shanghai, China) according to the manufacturer’s instructions. The eluted samples were tested by an anti-MBP or anti-HIS antibody (Beyotime, Shanghai, China). The primers used in this experiment are shown in Supplementary Table 1.

### RT-PCR and RT-qPCR

Total RNAs were extracted from Arabidopsis and NHCC plants using RNAprep Pure Plant Kit (TIANGEN, Beijing, China). The reverse transcription was carried out using the Hifair^®^Ⅲ One Step RT-qPCR SYBR Green Kit (YEASEN, Shanghai, China). RT-qPCR was performed with Hieff^®^qPCR SYBR Green Master Mix (YEASEN, Shanghai, China). The standard protocol of the kit was followed to mix the reaction components and perform the reactions (CFX Connect Real-Time PCR Detection System). 2^-ΔΔ^ CT method was used for calculating the relative expression level of the target gene (Livak and Schmittgen, 2001), and the specific primers used are listed in Supplemental Table 1.

### Yeast one hybrid (Y1H)

Y1H assay was conducted according to protocol (Clontech, cat. No. 630491). Briefly, the promoter fragment of *BcGSTF10*, containing GCC-box, was cloned from the genomic DNA of the NHCC. The amplified DNA fragments were ligated into the pAbAi vector. Bait reporter strains were created by integrating linearized pAbAi-BcGSTF10 plasmids into the Y1H Gold yeast strain, resulting in reporter strains. Next, pGADT7-BcCBF2 was fused into the pAbAi-BcGSTF10 bait strain, and validated on SD/-Ura/-Leu plates containing appropriate concentrations of AbA. The primers used in this experiment are shown in Supplementary Table 1.

### Electrophoretic mobility shift assay

The purified proteins of MBP-BcGSTF10, GST-BcCBF2 and His-BcICE1 were obtained as described in the pull-down assay experiment. Biotin-labeled and mutant probes were synthesized (Sangon Biotech, Shanghai, China). A Chemiluminescent EMSA Kit was used to perform the EMSA experiment (GS009, Beyotime, Shanghai, China). Briefly, DNA fragment (probe) was incubated with purified protein. Non-denatured acrylamide gel was used to separate the bound and free probes. Then, luminescent images were captured using a protein imaging system (Tanon 4600, Shanghai, China).

### Dual-luciferase assay

The Dual-luciferase assays were performed as described by Wang et al. (2022). In brief, the promoter sequences of *BcCBF2* and *BcGSTF10* were separately inserted into the pGreen0800-LUC vector to generate the reporter constructs. The effector constructs (*35S::BcICE1* and *35S::BcCBF2* were separately generated by recombining with the pGreenII 62-SK vector. The resultant vectors were then transformed into *A. tumefaciens* GV3101(psoup) (TOLOBIO, Shanghai, China). The Agrobacterium cells containing the reporter and effector construct were injected into tobacco leaves at a 1:10 ratio. Luminescence was detected with a live imaging apparatus. The activity of LUC and REN was measured by Dual-Luciferase Reporter Assay kit according to the instruction (Yeasen, Shanghai, China). All primers used in the dual-luciferase reporter assay are listed in Supplementary Table 1.

### Silencing of *BcGSTF10* in NHCC

VIGS assays were performed as described by Wang et al. (2022). Briefly, a 40-bp interference fragment of *BcGSTF10* and its antisense fragment were constructed into the *pTY* vector. Transform the constructed plasmid into NHCC seedlings through particle gun bombardment (Bio-Rad, PDS1000/He). The empty *pTY* plasmid was used as the control. Upon appearance of the mosaic phenotype, RNA extraction will be carried out on the susceptible leaves. qRT-PCR will then be performed to test its silencing efficiency, with the control used as a reference (Fig. 4B). The qRT-PCR-confirmed silenced lines will be used for subsequent experiments. Nucleotides used for VIGS assay are listed in Supplementary Table 2.

## RESULTS

### 1. BcICE1 interacts with BcGSTF10

In our previous study, we reported that BcICE1 plays a significant role in regulating cold stress in NHCC (Non-heading Chinese Cabbage). To gain a comprehensive understanding of the molecular mechanism associated with BcICE1 in NHCC, we used the Y2H system to identify protein that interact with BcICE1 and potentially modulate cold responses. Because both the full-length and the N-terminal region of BcICE1 activated the transcription of a GAL4-responsive reporter gene in yeast, we used the C-terminal region of BcICE1 (amino acid residues 254–446) as the bait. Sequence analysis of putative positive colonies suggested that a glutathione transferase gene interacts with BcICE1 in yeast, which we named *BcGSTF10*. To confirm the BcICE1–BcGTSF10 interaction, we introduced the full-length ICE1 into the Gal4 activation domain of the prey vector (AD-ICE1). The BD-BcGSTF10 and AD-ICE1 plasmids were co-transformed into yeast to examine protein–protein interactions. To more precisely identify the BcICE1 region responsible for the interaction with BcGSTF10, we fused four truncated BcICE1 variants to the Gal4 activation domain of the prey vector and full length BcGSTF10 into the bait vector. We found that only BcICE1-F2 and BcICE1-F4 are capable of interacting with BcGSTF10. This implies that the bHLH region of ICE1 is not necessary for the interaction with BcGSTF10, but rather the ACT region (Fig. 1A, B). To further confirm the interaction between BcGSTF10 and BcICE1, bimolecular fluorescence complementation (BiFC) assays were conducted. Strong yellow fluorescence signal was observed in the nuclei of cells co-expressing YFP^n^-BcICE1 and YFP^c^-BcICE1 (Fig. 1C). On the contrary, no fluorescence was detected in negative controls in which BcICE1-YFP^n^ was co-expressed with YFP^c^ empty vector or BcGSTF10-YFP ^c^ was co-expressed with YFP^n^ empty vector. In addition, in vitro pull-down assay results showed MBP-BcGSTF10 was pulled down by His-BcICE1 (Fig. 1D). Taken together, these results demonstrate that BcICE1 physically interacts with BcGSTF10 in plant cell nuclei, suggesting that BcGSTF10 functions as an interacting partner of BcICE1, potentially involved in regulating cold stress in NHCC

**Figure1.**
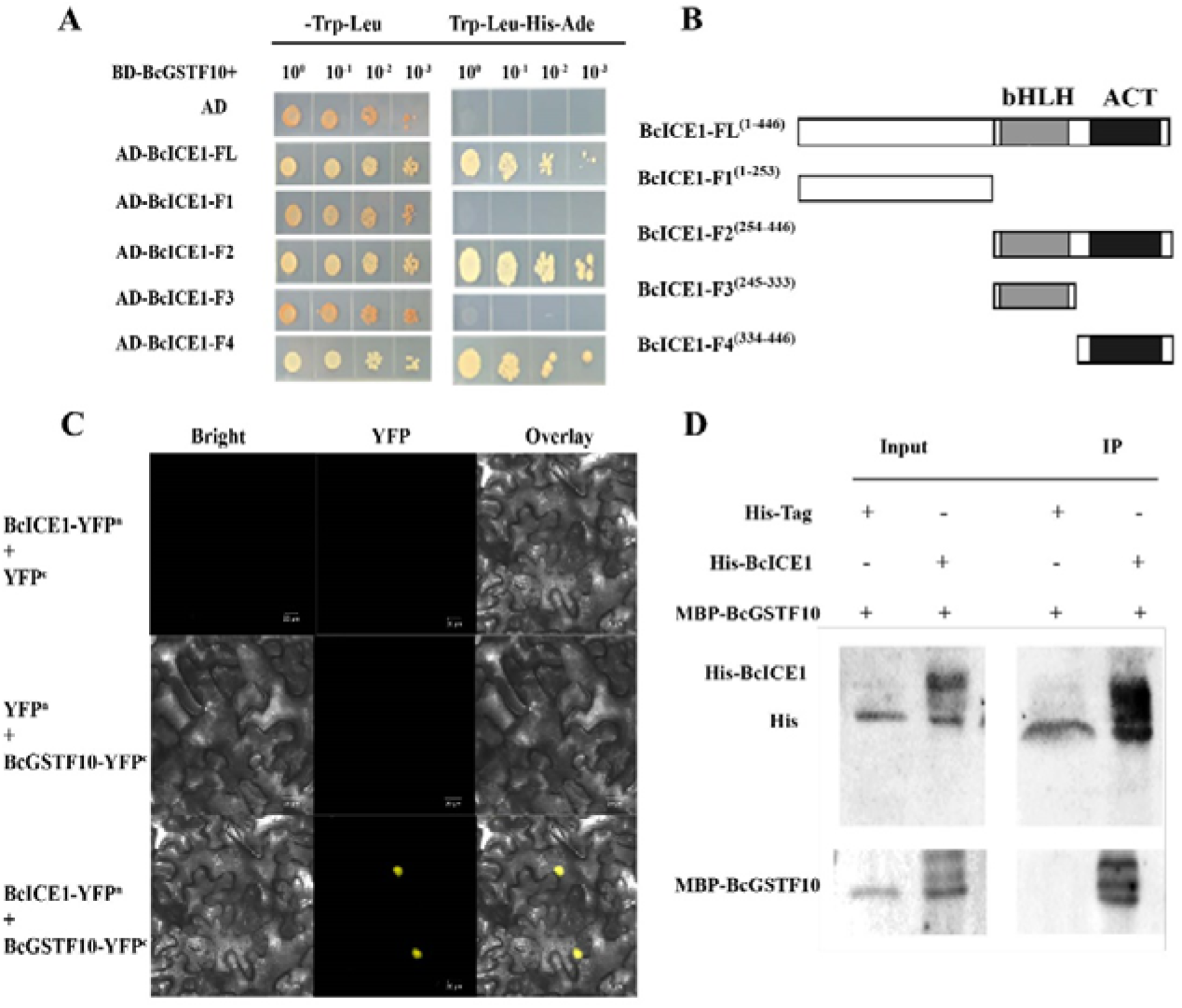
BcGSTF10 interact with BcICE1. (A, B) BcSGTF10 interact with BcICE1 in a yeast two-hybrid assay, and BcICE1-F3 fragment is required for this interaction. The empty vector pGADT7 was used as negative control. (C) BcSGTF10 interact with BcICE1 in a bimolecular fluorescence complementation (BIFC) assay. Bars=20μm. (D) BcGSTF10 interact with BcICE1 in a pull-down assay. MBP-tagged BcGSTF10 incubated with His-BcICE1 or His protein, and the immunoprecipitated proteins was detected with anti-his antibody.

### 2. BcGSTF10 Positively Regulates cold stress

*BcGSTF10* comprised 3 exons and 2 introns (Supplementary Fig. 1A). Its 648-base pairs (bp) open reading frame (ORF) encoded a predicted protein of 215 amino acids (aa). Amino acid sequence alignment demonstrated that BcGSTF10 contained two conserved GST domain (Supplementary Fig. 1A). Phylogenetic analysis suggests that the BcGSTF10 proteins were closely related to GSTF10 from *Brassica napa* and *Raphanus sativus* (Supplementary Fig. 1B). To explore the position and potential function of BcSTF10, the gene was fused with a green fluorescent protein (GFP) reporter gene, and *35S:BcGSTF10-GFP* was transiently expressed in *Nicotiana tabacum*. Results showed that BcGSTF10 was localized in the nucleus and cell membrane (Supplementary Fig. 1C). Further, we examine the expression pattern of *BcGSTF10* under low-temperature stress treatment by RT-qPCR. It can be seen from the trend of the curve that *BcGSTF10* could be slight upward in a short time. However, when exposed to sustain low temperature the mRNA level significantly reduced (Supplementary Fig. 1D).

To investigate the role of BcGSTF10 in cold stress, *BcGSTF10* transgenic NHCC plants were successfully obtained through tissue culture-mediated transformation (Supplementary Fig2. A, B). Freezing treatment experiments were conducted using transgenic BcGSTF10 plants in NHCC (Fig.2A, B). The transgenic *BcGSTF10* plants exhibited significantly enhanced freezing tolerance and lower relative electrolyte leakage compared to the control. Meanwhile, the 35S-promoter-driven *BcGSTF10-GFP* and GFP empty vector were expressed in NHCC leaves via Agrobacterium-mediated transient transformation. After cold stress treatment, the results revealed that DAB and NBT staining were shallower in leaves of overexpressing *BcGSTF10* plants compared with that for empty vector control (Fig. 2C). In addition, relative ion leakage demonstrated that empty vector control had more severe cell membrane damages (Fig. 2D). qRT-PCR results showed that overexpression of *BcGSTF10* promoted the expression of *BcCBFs* and *CBFs* target genes, including *BcCOR15A* and *BcCOR47* in NHCC. These results demonstrate that *BcGSTF10* could positively regulate freezing tolerance in NHCC.

**Figure 2.**
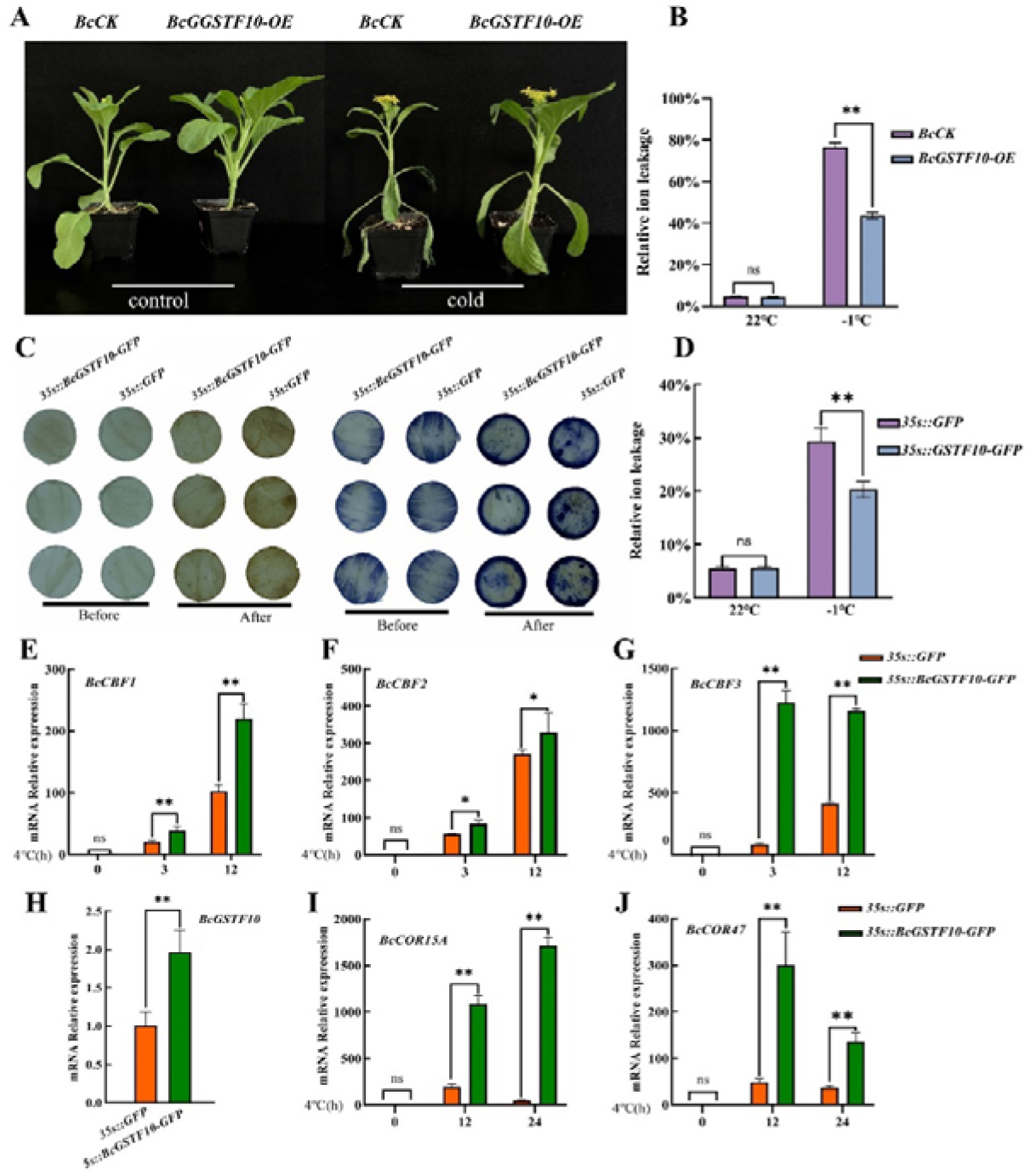
Cold phenotype of *BcGSTF10* transgenic plants or leaves in vitro. (A-B) cold tolerance assay in *BcGSTF10* transgenic NHHCC. (A) Phenotypes of *BcCK* and *BcGSTF10* overexpression plants after freezing treatment. Photographs show representative plants after recovery. Plants grown for 25 days under 22LJ conditions were transferred to -1LJ for 12 hours of treatment. (B) Relative ion leakage of plants after the freezing assays described in (A). (C-J) cold tolerance assay using *BcGSTF10* overexpressing leaves discs. (C) DAB and NBT staining under freezing stress in control and *BcGSTF10* overexpression leaves discs. (D) Relative ion leakage under freezing stress in control and *BcGSTF10* overexpression leaves discs. (E-J) Expression of endogenous *BcCBFs* (E-G), *BcGSTF10* (H) and *BcCORs* (I-J) genes in *BcGSTF10* overexpression leaves discs under cold stress. The data are the mean values of three independent experiments ± SD. **P < 0.01 (Student’s t-test)

Moreover, *BcGSTF10* was heterologously expressed in *Arabidopsis*, the transgenic plants overexpressing BcGSTF10 driven by a super promoter displayed freezing tolerance phenotype compared with wild-type plants (Fig. 3A). Under non-acclimated conditions, approximately 55% of the transgenic lines survived after freezing treatment at -5°C for 1 h, whereas only 25% wild type survived (Fig. 3B). Next, electrolyte leakage and ROS accumulation were examined between wild and overexpression line. No obvious difference was observed for the electrolyte leakage and ROS accumulation between the wild-type and *BcGSTF10* overexpressing plants under normal condition. However, when exposed to chilling environment the electrolyte leakage and ROS accumulation in wild lines were remarkably increased compared to transgenic lines (Fig. 3C-E). Consistently, the expression of *CBFs* (*CBF1*, *CBF2*, *CBF3*) and downstream genes *CORs*, including *COR15B*, *RD29A* and *KIN1*, was strongly upregulated in *BcGSTF10* overexpression transgenic *Arabidopsis* compare to the wild lines (Fig. 3F-K). These results demonstrate that *BcGSTF10* enhances plant cold tolerance under cold stress conditions by promoting the transcription of *CBFs* and their target genes.

**Fig3.**
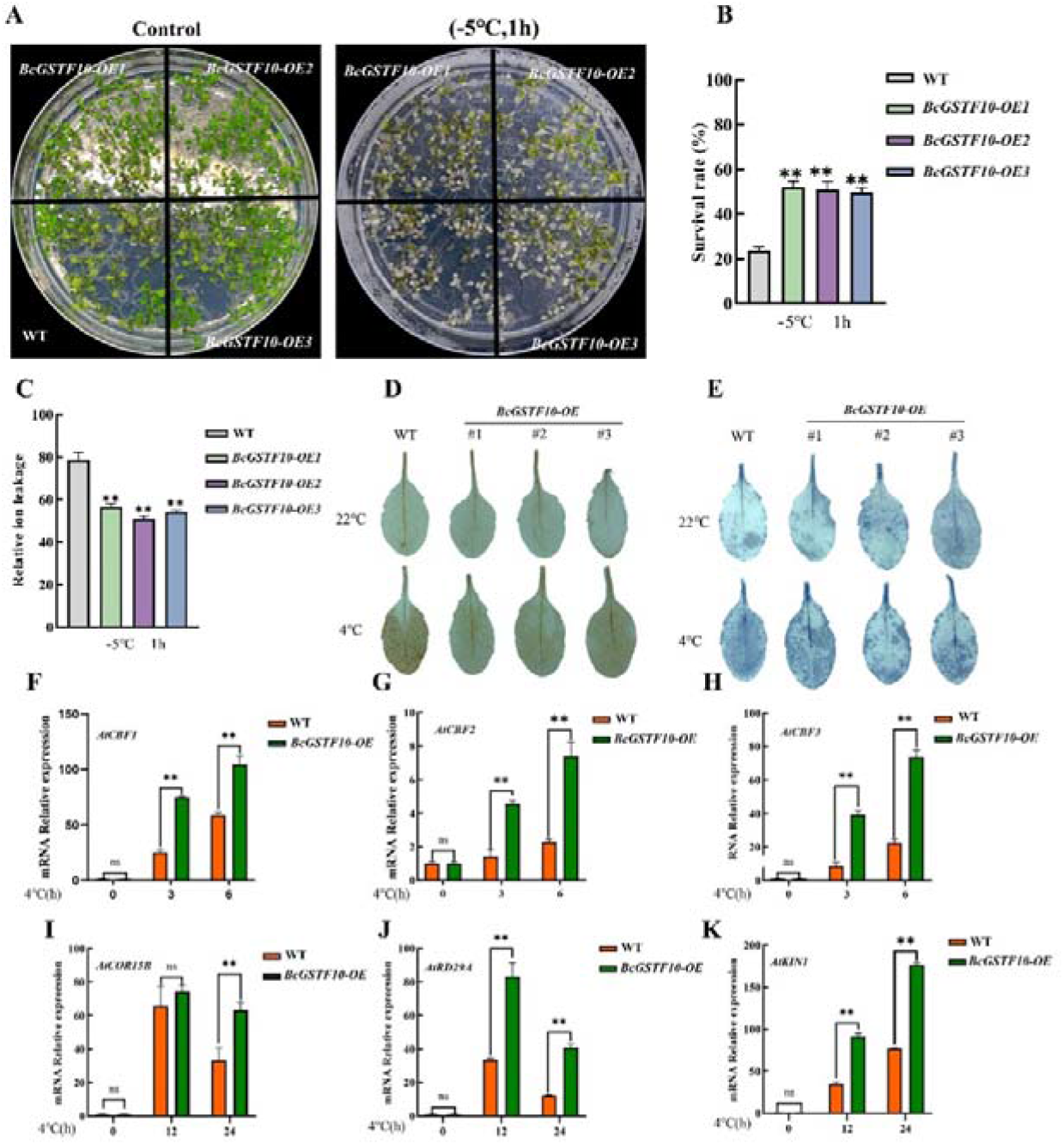
Cold-stress phenotypes of *BcGSTF10* transgenic *Arabidopsis* seedlings. (A) Phenotypes of WT and *BcGSTF10* overexpression plants after freezing treatment. The plants were grown on ½ MS plates at 22°C for 2weeks then a −5°C treatment for1h. (B) Survival rates of plants after the freezing assays described in (A) (C) Relative ion leakage of plants after the freezing assays described in (A) (D-E) DAB (D) and NBT (E) staining under cold stress. Plant leaves were treated at 4 °C for 3 d, and then leaves were dyed with DAB and NBT. (F-K) Expression of *CBFs*(C) and *CORs*(D) in WT and *BcGSTF10* transgenic plants under cold stress. Data are the means of three independent experiments ± SD. *P < 0.05, **P < 0.01 (Student’s t-test).

### 3. Silencing of *BcGSTF10* reduced cold tolerance in NHCC plants

To further confirm the function of *BcGSTF10* in cold stress, a VIGS assay was performed to examine the effect of *BcGSTF10* silencing in NHCC under cold stress. After treatment at -1°C for 24 h, the *pTY-BcGSTF10* lines showed substantially reduced cold tolerance compared to the wild type (Fig. 4A, B). DAB and NBT staining indicate that *pTY-BcGSTF10* plants exhibit increased production of H_2_O_2_ and O^2-^ compared to *pTY* lines following cold stress (Fig. 4C, D). Relative ion leakage suggests that *pTY-BcGSTF10* plants exhibit more severe cell membrane damage after cold stress compared to *pTY* plants (Fig. 4E). Additionally, the expression of *CBFs* and downstream *CORs* were strongly reduced in *pTY-BcGSTF10* lines compared to *pTY* lines (Fig. 4F-J). Based on these results, we conclude that *BcGSTF10* is crucial for cold tolerance in NHCC.

**Fig4.**
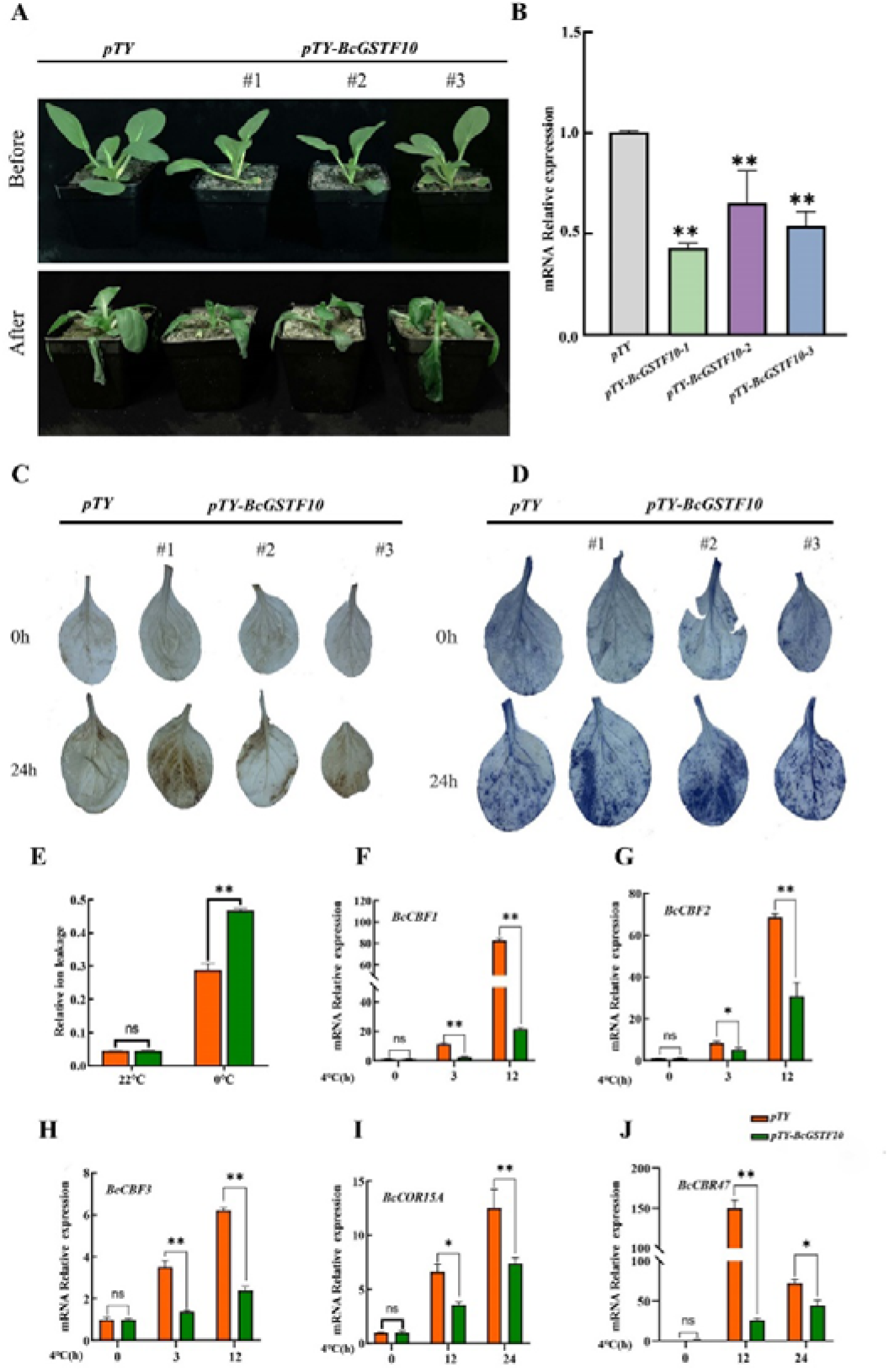
Cold-stress phenotypes of *BcGSTF10* silenced in NHCC. (A) Phenotypes of *pTY* and *pTY-BcGSTF10* plants after freezing treatment. The silenced NHCC growth in 22LJ for 10 days and then a -1LJ treatment for 24h. Photographs show representative plants after recovery. (B) The expression pattern of *BcGSTF10* in *pTY* and *pTY-BcGSTF10* in NHCC. (C-D) DAB (C) and NBT (D) staining under cold stress. Silenced NHCC were treated at 0 °C for 3 d, and then leaves were dyed with DAB and NBT. (E) Relative ion leakage of plants after the freezing assays described in (C-D) (F-J) Expression of CBFs(C) and CORs(D) in WT and BcGSTF10 transgenic plants under cold stress. Data are the means of three independent experiments ± SD. *P < 0.05, **P < 0.01 (Student’s t-test).

### 4. BcGSTF10 enhances the transcriptional activity of BcICE1

In *Arabidopsis*, ICE1 regulates cold-stress response by promote the expression of *CBFs* when exposed to cold stress (Chinnusamy et al., 2003). In this study, a G-box was found in *BcCBF2* promoter, in order to investigate whether *BcCBF2* are direct targets of BcICE1, we performed an electrophoretic mobility shift assay (EMSA). EMSA results showed that BcICE1 directly bound to a G-box site (CACGTG) of the *BcCBF2* promoter in vitro (Fig. 5A, B). Considering that BcGSTF10 acts as an interaction partner of BcICE1 and also modulates cold stress, we wondered whether transcriptional activation of *BcICE1* genes is modulated by BcGSTF10. EMSA assay showed that with the addition of BcGSTF10 protein, the binding strength of BcICE1 to the promoter of *BcCBF2* increased, suggesting that BcGSTF10 promotes the binding of BcICE1 to the promoter of *BcCBF2* (Fig. 5C). In addition, dual-luciferase assays were performed to assess the effect of BcGSTF10 on the transcriptional activation of BcICE1 (Fig. 5D). The fluorescence assay results showed that the expression of BcCBF2 was significantly enhanced when BcICE1 and BcGSTF10 co-expressed (Fig. 5E, F). These results indicate that BcGSTF10 activate BcCBF2 expression through enhances the transcriptional activation activity of BcICE1.

**Fig5.**
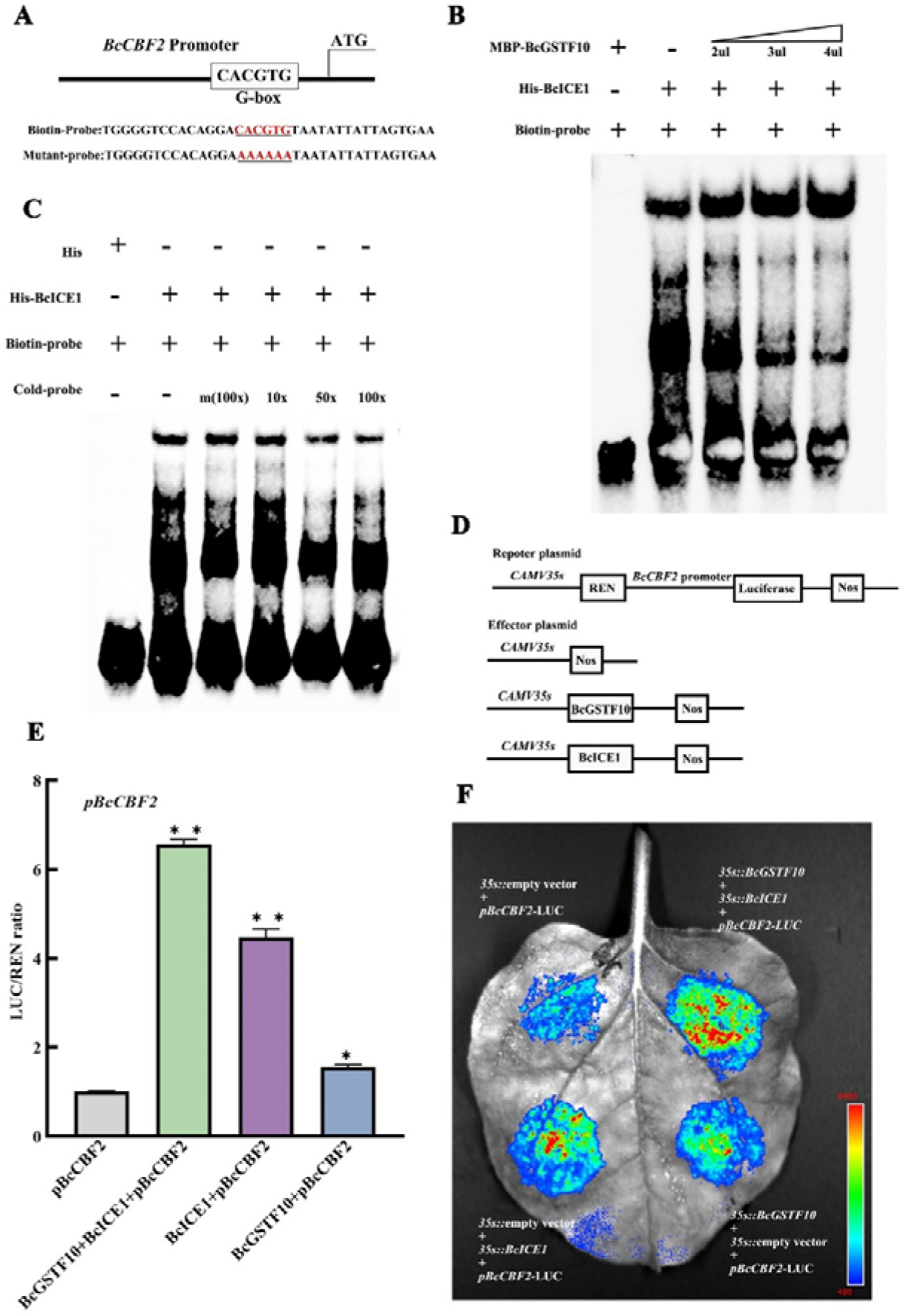
BcGSTF10 enhances the transcriptional function of BcICE1. (A) Schematic diagram showing the *BcCBF2* promoter probe used for EMSAs. The BcICE1 binding site (G-box) is indicated. (B) EMSAs showing that BcGSTF10 enhances the binding ability of BcICE1 to the *BcCBF2* promoter. (C) EMSAs showing that BcICE1 binding to the *BcCBF2* promoter. (D) Schematic representation of the LUC reporter vector containing the BcCBF2 promoter and the effector vectors expressing BcGSTF10 or BcICE1under the control of the 35S promoter. (E) LUC/REN ratio detected by the reporter system described in (D), testing the effects of BcGSTF10, BcICE1, and BcGSTF10+BcICE1 on the expression of BcCBF2. (F) Luciferase imaging assay showing that BcGSTF10 enhances the transcriptional activity in *N. benthamiana* leaves. The fluorescence intensity of the control was used as the reference. Data are the means of three independent experiments ± SD. *P < 0.05, **P < 0.01 (Student’s t-test).

### 5. BcGSTF10 Acts Directly Downstream of BcCBF2

In order to figure out the role of *BcGSTF10* in long term cold stress, promoter sequences were analyzed. Specific binding sites were found for the promoter fragment of *BcGSTF10* containing the CRT/DRE cis-element (Fig. 6A). To further dissect whether BcCBF2 can directly regulate BcGSTF10. A Y1H yeast assay was conducted, the promoter of *BcGSTF10* fused to pAbAi vector, as well as BcCBF2 with the GAL4-activating domain (AD). Y1H assay suggesting that co-expressing pABAi-BcGSTF10 and AD-BcCBF2 were able to grow on the SD/−Leu/−Trp/−His medium supplemented with 200 ng ml^-1^ AbA after a 1000-fold dilution in the bacterial solution, while no yeast cells transformed with an empty vector were found to grow (Fig. 6C). Further, EMSA showed that BcCBF2 directly bound the CRT/DRE cis-element of the *BcGSTF10* promoter. Unlabeled competitive probes reduced this binding activity and mutant probe lost binding activity (Fig. 6A, B). As a means of illustrating how *BcGSTF10* was regulated by BcCBF2, we carried out dual-luciferase assays. The *BcCBF2* was driven by 35S-promoter, and the promoter of *BcGSTF10* were cloned into pGreenII 0800-LUC.We found that a drastic reduction fluorescence of overexpression BcCBF2 compared with empty vector control (Fig. 6D-F). All of this demonstrates that *BcGSTF10* was directly inhibited by BcCBF2.This appears to account decreased gene expression of *BcGSTF10* when exposed continuously in cold stress (Supplementary Fig. 1D).

**Fig6.**
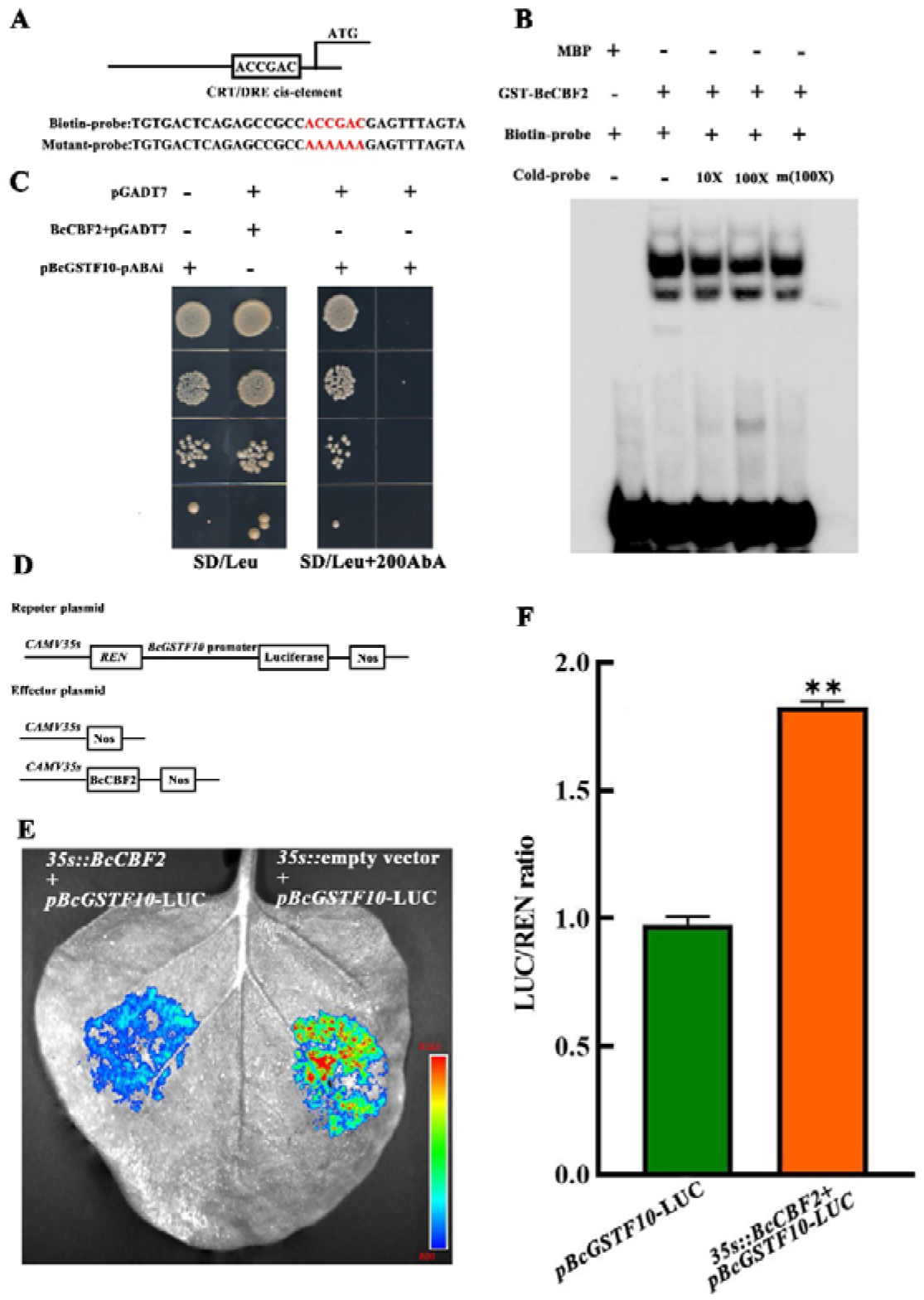
BcCBF2 directly bind to the promoter of *BcGSTF10*. (A) Schematic diagram showing the *BcGSTF10* promoter probe used for EMSAs. The BcCBF2 binding site (CRT/DRE cis-element) is indicated. (B) EMSAs showing that BcCBF2 binding to the *BcGSTF10* promoter. (C) BcCBF2 binding to the *BcGSTF10* promoter in a yeast two-hybrid assay. (D) Schematic representation of the LUC reporter vector containing the *BcGSTF10* promoter and the effector vectors expressing BcCBF2 under the control of the 35S promoter. (E) Luciferase imaging assay showing that BcCBF2 binding to the *BcGSTF10* promoter. (F) LUC/REN ratio detected by the reporter system described in (D). testing the effects of BcCBF2 on the expression of BcGSTF10. The fluorescence intensity of the control was used as the reference. Data are the means of three independent experiments ± SD. *P <0.05, **P < 0.01 (Student’s t-test).

### 6. BcGSTF10 Regulates plant Growth

In addition to cold tolerance, *BcGSTF10* gene may participate in plant growth. Overexpression of *BcGSTF10* leads to increased plant size and leaf number in both NHCC and *Arabidopsis* (Fig. 2A, 7A; Supplementary Fig.3A-C). Compared to the control, the growth of the *BcGSTF10*-silenced plants are significantly inhibited, exhibiting symptoms of dwarfism with smaller size and fewer leaves (Fig. 7B; Supplementary Fig.3D-E). In *Arabidopsis*, the cross sections of the first stem internodes of transgenic and wild-type (WT) plants were compared. In transgenic plants, the area of the cross-section was significantly larger than in the WT plants. Subsequent observations indicated that cell size did not significantly differ between the two groups, but cell number significantly increased in transgenic plants (Fig. 7C). We conducted a comparative analysis of shoot apical meristem cells between *pTY-BcGSF10* lines and *pTY* lines, and found that the latter exhibited a higher density of stem cells in the apical meristem region (Fig. 7D). These results suggest that *BcGSTF10* may promote growth by influencing cell division. Further, RT-qPCR reveal that growth-related genes *BcRI1/AtBRI1*, *BcDWF4/AtDWF4*, *BcYUCCA8/AtYUCCA8*, were up-regulated compared with the wild type, while these genes were down-regulated in *pTY-BcGSTF10* plants (Fig. 7F-I). In addition, studies have shown that overexpression of *CBF2* in *Arabidopsis* results in a dwarf phenotype (Gilmour et al, 2004; Zhao et al 2016). In order to determine whether the *BcCBF2* gene has a similar function to that in *Arabidopsis*, we performed a heterologous overexpression experiment of the *BcCBF2* gene in Arabidopsis (Supplementary Fig.2D). The results showed that *BcCBF2* overexpressing plants phenotypically resembled Arabidopsis *AtCBF2*, whereas overexpression of *BcCBF2* gave rise to an opposite phenotype when compared to *BcGSTF10* (Fig. 7E). Next, we conducted a transgenic experiment by crossing plants that overexpressed *BcCBF2* and *BcGSTF10* genes to generate *BcGSTF10-OE BcCBF2-OE* plants. (Supplementary Fig. 2E). The results showed that *BcGSTF10-OE BcCBF2-OE* plants can reverse the dwarf phenotype caused by the overexpression of *BcCBF2* gene (Fig. 7E). These findings imply that the influence of *BcCBF2* on the growth and development of NHCC could potentially be mediated through the modulation of *BcGSTF10* levels.

**Fig7.**
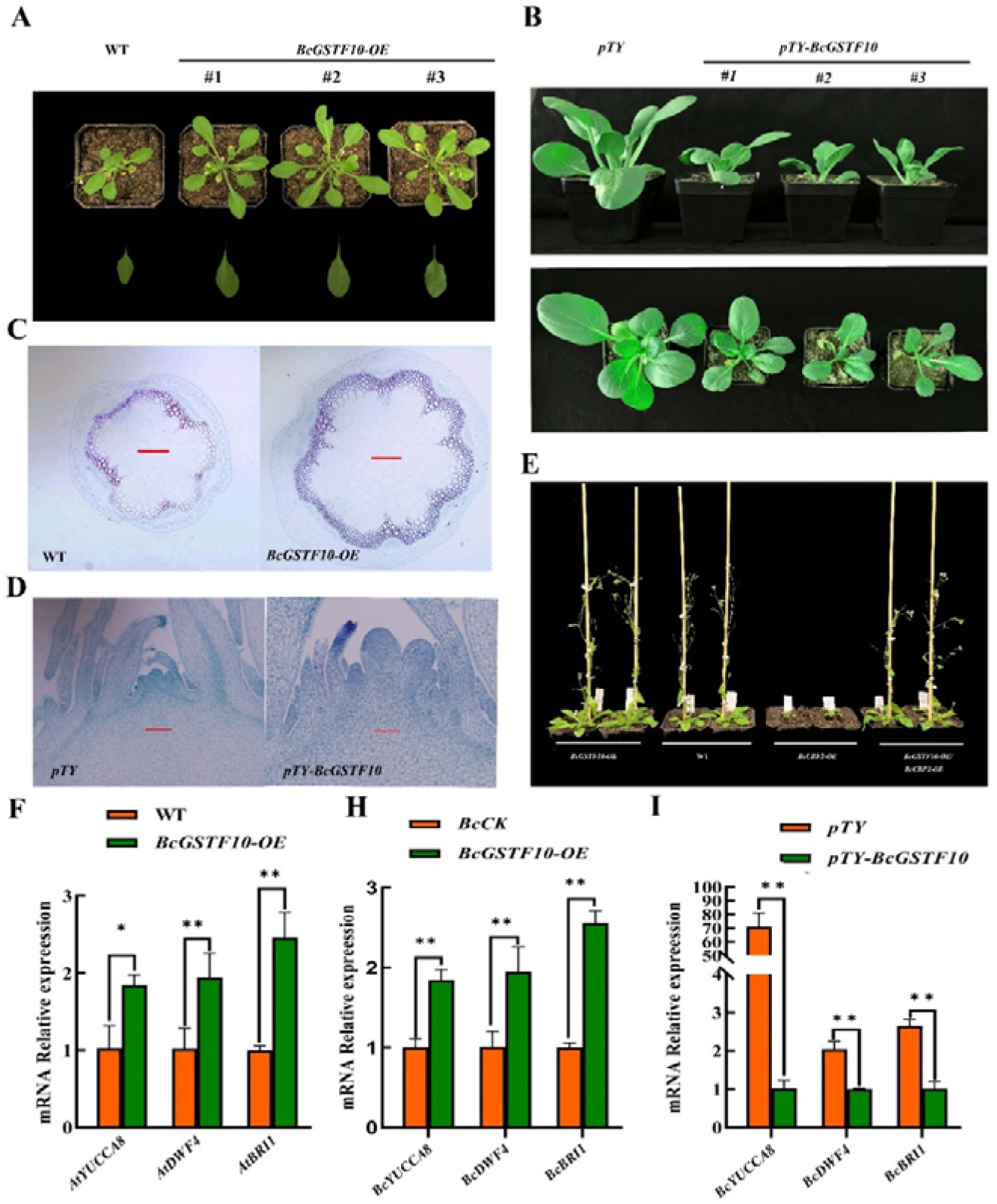
BcGSTF10 affects the growth of plants. (A) The phenotype of WT and *BcGSTF10* overexpressing Arabidopsis after 35 days of growth in a light incubator. (B) The phenotype of *pTY* and *pTY-BcGSTF10* plants after silenced 3 weeks. (C) The cross section of the first internode stem tissues in WT and BcGSTF10 overexpression *Arabidopsis*. (D) The cross section of shoot apical meristem in *pTY* and *pTY-BcGSTF10* plants. (E) The phenotype of *BcGSTF10-OE*, WT, *BcCBF2-OE* and *BcCBF2-OE/ BcGSTF10-OE Arabidopsis* after 45 days of growth in a light incubator. (F) qRT-PCR analysis the expression level of *AtYUCCA8*, *AtDWF4*, *AtBRI1* in WT and overexpression *BcGSTF10 Arabidopsis*. The value for WT was set to 1. (H) qRT-PCR analysis the expression level of *BcYUCCA8*, *BcDWF4*, *BcBRI1* in control and overexpression *BcGSTF10* NHCC. The value for control line was set to 1. (I) qRT-PCR analysis the expression of *BcYUCCA8*, *BcDWF4*, *BcBRI1* in *pTY* and *pTY-BcGSTF10* NHCC. The value for *pTY* line was set to 1.

## Discussion

Cold stress is one of the major abiotic factors that limit plant growth, development, and economic crop yield in certain regions (Ding et al., 2022). Cold stress stimulates the accumulation of reactive oxygen species (ROS) in plant tissues (Lee et al., 2002; Dong et al., 2009; Cen et al., 2018; Shi et al., 2019). The accumulating ROS cause oxidative stress, which damages the plant cell membrane, decreases enzyme activity, and inhibits the rate of photosynthesis and protein translation (Huner et al, 2012). Plants use GSTs (Glutathione S-transferase) to catalyze the combination of harmful xenobiotics or oxidized products with GSH (Redueed glutathione), facilitating the metabolism and clearance of these chemicals (Ahmad et al., 2016).GSTs have been identified to be critical players in managing biotic and abiotic stress, promoting plant growth and development, regulating phytohormone and cell signaling, maintaining redox homeostasis, controlling the biosynthesis and transport of secondary metabolites such as anthocyanin, as well as mediating cell death in plants (Nianiou-Obeidat et al. 2017; Chen et al. 2007; Singhal et al. 2015; Dixon et al. 2010 and Hu et al. 2016). In our research, we discovered a glutathione S-transferase, *BcGSTF10*, which play a positive role in regulating cold tolerance in both NHCC and *Arabidopsis* (Fig. 2A-J; Fig. 3A-K). Consistent with this result, the *BcGSTF10* silenced lines exhibited a pronounced cold-sensitive phenotype in NHCC, as compared to the control. (Fig. 4A-J). Early studies in *Arabidopsis thaliana* have shown that the expression of the *BcGSTF10* gene is induced by drought stress rather than cold stress. Therefore, it is inferred that *BcGSTF10* may play a role in the response to drought stress rather than in response to low-temperature stress (Kiyosue et al.,1993). However, research conducted on NHCC has shown that *BcGSTF10* expression can be up-regulated at low temperatures within a short period of time, but prolonged exposure to low temperatures inhibits its expression (Supplementary Fig.1A).

Additionally, our screening and subsequent in vivo/in vitro experiments have all confirmed the interaction between BcGSTF10 and BcICE1. ICE1 is a key positive regulatory transcription factor in the low-temperature signaling pathway of plants, and its protein activity is subject to multiple post-transcriptional modifications (Chinnusamy et al., 2003; Miura et al., 2007; Ding et al., 2015; Li et al., 2017a; Ye et al., 2019). However, studies have limited information on the interactions between ICE1 and other factors, as well as the roles played by such interactions. Evidence from previous studies indicated that ICE1 bind to multiple MYC recognition sites on the promoters of CBF genes (Chinnusamy et al., 2003). Our research suggests that BcICE1 is capable of binding to the G-box region in the promoter of BcCBF2 (Fig. 5A, C). In addition, the interaction between BcGSTF10 and BcICE1 can impact the binding activity of BcICE1 with BcCBF2 promoter (Fig. 5B, D-F). Furthermore, it was discovered that the promoter region of BcGSTF10 has a CRT/DRE cis-element element (Fig. 6A). The CBF proteins are capable of recognizing a cis-element known as CRT/DRE, which encompasses the conserved CCGAC sequence (Stockinger et al., 1997; Liu et al., 1998; Gilmour et al., 1998). Through Y1H, electrophoretic mobility shift assay (EMSA), and luciferase assay, it was found that BcCBF2 directly binds to the promoter of *BcGSTF10* and negatively regulated the transcription of *BcGSTF10* (Fig. 6A-F). These findings may shed light on the mechanism behind the reduced expression of the BcGSTF10 gene during prolonged exposure to low temperatures (Supplementary Fig. 1A).

Evidence suggests that auxins, cytokinin, salicylic acid, methyl jasmonate, ethylene, and other hormones induce the expression of GSTs, highlighting the crucial role of GSTs in the growth and development of plants (Gong et al., 2005; Moons et al., 2005; Ahmad L et al., 2016). Previous reports have demonstrated that *AtGSTU17* is involved in various developmental processes such as seedling growth, elongation of the hypocotyl, accumulation of anthocyanins, and far-red light-mediated inhibition of greening (Jiang et al., 2010). Our research findings demonstrate that the *BcGSTF10* gene exerts a positive influence on the regulation of plant growth. Through overexpression of *BcGSTF10*, we observed a significant increase in plant size and rosette leaves number (Fig. 7A; Supplementary Fig. 3A-C). Conversely, we observed a notable growth inhibition phenotype in silenced lines (Fig. 7B; Supplementary Fig. 3D-F). Paraffin sections suggest that these changes may be associated with alterations in cell division (Fig. 7C, D). The *CBFs* genes have been regarded as the most extensively studied genetic factors involved in plant response to cold stress, serving as a crucial regulator in coordinating various adaptive mechanisms (Agarwal et al., 2006; Chinnusamy et al., 2003; Doherty et al., 2009; Shi et al., 2012). Heterologous overexpression of *BcCBF2* in *Arabidopsis thaliana* resulted in transgenic plants with a stunted growth phenotype (Fig. 7E), consistent with previous findings on *CBFs* in *Arabidopsis thaliana* (Achard et al., 2008; Gilmour et al., 2000; Gilmour et al., 2004). Furthermore, we found that the *BcCBF2-OE BcGSTF10-OE* plants were able to rescue the dwarf phenotype of BcCBF2 (Fig. 7E). This indicates that the growth retardation caused by overexpression of *BcCBF2* may be caused by the inhibition of *BcGSTF10* expression.

Based on previous findings and our evidence, we propose a model in which *BcGSTF10* balances between plant growth and cold stress (Fig. 8). Non-heading Chinese cabbage displays steady levels of *BcGSTF10* expression and normal growth in optimal temperature conditions. In short-term cold stress, cold signals activate *BcGSTF10* expression, promoting the expression of downstream cold-related signaling genes via BcICE1 activation and facilitating resistance to low temperature. However, prolonged exposure to low temperature increases the accumulation of BcCBF2 protein, leading to *BcGSTF10* expression inhibition and, subsequently, inhibited growth.

**Fig8.**
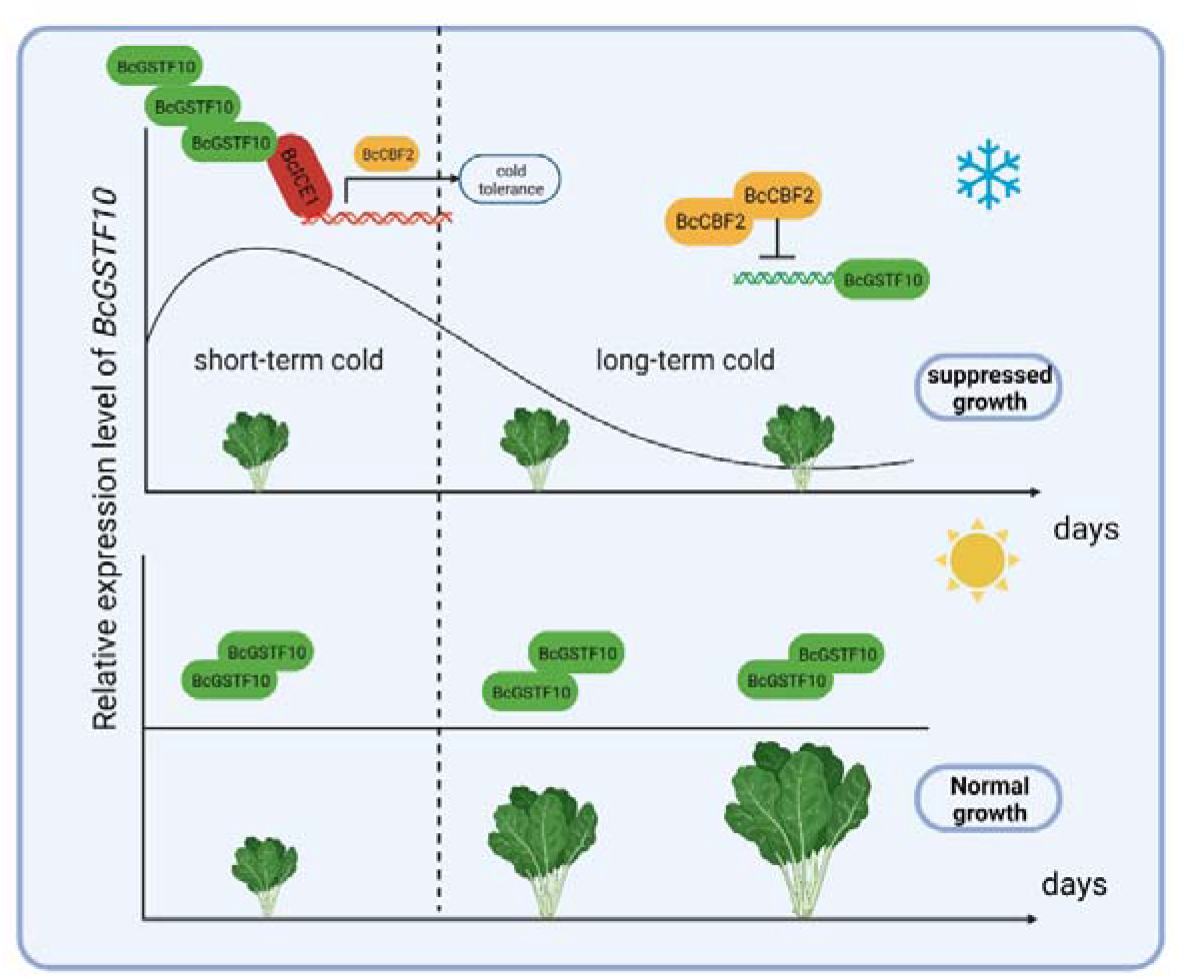
Model for *BcGSTF10* effect cold stress and growth in NHCC. The expression of *BcGSTF10* remains at a steady level, enabling normal plant growth, under optimal thermal conditions. During brief exposure to cold stress, the ability of BcGSTF10 to perceive low temperatures activates ICE1 transcription, conferring enhanced cold tolerance. However, prolonged exposure to cold stress results in a substantive accumulation of BcCBF2, which interrupts BcGSTF10 expression and inhibits growth.

## Supplementary data

**Supplemental Table 1.** Primers used for gene expression and vector construction.

**Supplemental Table 2.** The CDs sequence of *BcGSTF10*.

**Supplemental Figure 1.** The expression and structure analysis of BcGSTF10.

**Supplemental Figure 2.** Identification of overexpressing transgenic lines in NHCC and *Arabidopsis*.

**Supplemental Figure 3.** B*c*GSTF10 contributing to increased biomass and yield.

## Author contributions

Y.L and X.L.H designed the research. S.Y.L H., Y. W and L.Y.R performed the research J.J.R and G.P.W analyzed the data. Y.L.S wrote the manuscript. Q.D and Y.L revised the article.

## Data availability

The data that supports the findings of this study are available in the Supporting Information of this article.

